# Microbial subnetworks related to short-term diel O_2_ fluxes within geochemically distinct freshwater wetlands

**DOI:** 10.1101/363226

**Authors:** Dean J. Horton, Matthew J. Cooper, Anthony J. Wing, Peter S. Kourtev, Donald G. Uzarski, Deric R. Learman

## Abstract

O_2_ concentrations often fluctuate over diel timescales within wetlands, driven by temperature, sunlight, photosynthesis, and respiration. These daily fluxes have been shown to impact biogeochemical transformations (e.g. denitrification), which are mediated by the residing microbial community. However, little is known about how resident microbial communities respond to diel dramatic physical and chemical fluxes in freshwater wetland ecosystems. In this study, total microbial (bacterial and archaeal) community structure was significantly related to diel time points in just one out of four distinct freshwater wetlands sampled. This suggests that daily environmental shifts may influence wetlands differentially based upon the resident microbial community and specific physical and chemical conditions of a freshwater wetland. However, when exploring at finer resolutions of the microbial communities within each wetland, subcommunities within two wetlands were found to correspond to fluctuating O_2_ levels. Microbial taxa that were found to be susceptible to fluctuating O_2_ levels within these subnetworks may have intimate ties to metabolism and/or diel redox cycles. This study highlights that freshwater wetland microbial communities are often stable in community structure when confronted with short-term O_2_ fluxes, however, specialist taxa may be sensitive to these same fluxes.

## INTRODUCTION

Diel O_2_ fluctuations have been observed within aquatic systems, including freshwater wetlands, with near anoxic levels occurring at night and elevated O_2_ concentrations occurring during daytime hours (Cornell & Klarer, 2008; Reeder, 2011; Maynard et al., 2012). In aquatic ecosystems other than wetlands (such as oceans and salt marshes), microbial community activity and structure can respond to short-term daily changes in environmental conditions (Ghiglione et al., 2007; Ottesen et al., 2014; Andrade et al., 2015; Morris et al., 2016; Kearns et al., 2017). As a consequence, geochemical processes can respond daily to diel environmental fluxes (Jørgensen et al., 1979; Laursen & Seitzinger, 2004; Harrison et al., 2005), as many geochemical transformations are driven by microbial communities that are sensitive to shifting environmental variables such as O_2_, pH, temperature, and light levels. Diel O_2_ fluxes are of particular interest within freshwater wetlands, as oxygen availability can influence microbial community metabolism, and thereby influence important transformations of redox-sensitive elements which commonly occur within wetlands, including (but not limited to) denitrification, methanogenesis, and oxidation and reduction of iron, manganese, and sulfate (Firestone & Davidson, 1989; Ehrlich, 1997; Beck & Bruland, 2000; Venterink et al., 2003; Reddy & DeLaune, 2008). Furthermore, shifts in microbial metabolic processes can directly control rates of carbon mineralization (Thomas et al., 1996; Kuehn et al., 2004), and in turn, regulate greenhouse gas emission and pollution mitigation in freshwater wetlands.

While evidence suggests that diel fluxes of O_2_ can influence microbial communities in several aquatic systems, the effects of environmental diel fluxes on microbial community structure and activity within freshwater wetlands has remained understudied. Specifically, current research on microbial community response to diel fluxes in freshwater wetlands has been limited to plant rhizosphere microbial communities. Rhizosphere community structure (i.e., beta diversity) has been consistently found to be stable throughout diel fluxes, while expression of functional genes appears to be idiosyncratic in response to diel fluxes based upon the functional community targeted. Xu et al. (2012) found that microbial communities were stable within rice plant rhizospheres, while *mcrA* (methyl-coenzyme M reductase) gene expression responded to diel fluxes, suggesting that methanogen activity may specifically respond to daily O_2_ fluctuations. In another study, both total microbial community structure, ammonium-oxidizer community structure, and *amoA* expression remained largely stable within wetland plant rhizospheres (Nikolausz et al., 2008). These studies suggest that subsets of microbial communities, possibly composed of microorganisms with redox-sensitive metabolisms, may respond uniquely to environmental diel fluxes. While insights have been gained on microbial community response to diel fluxes in studies of wetland rhizospheres, it is necessary to examine the influence of diel fluxes on microbial communities in the water column of freshwater wetlands. Recent research in a salt marsh has highlighted that active soil microbial communities remain stable in the face of diel environmental fluxes, while active microbial communities shift with diel fluxes within the water column (Kearns et al., 2017).

In this study, we sought to explore 1.) whether microbial community structure, or subnetworks of microbial taxa, varied between dawn and dusk time points across four distinct freshwater wetlands, and 2.) whether existing variability in microbial communities was related to environmental fluxes (e.g., O_2_, pH, and temperature). Throughout the summers of 2015 and 2016, four geochemically distinct freshwater wetlands within Michigan, U.S.A. were sampled at dawn and dusk time points for two consecutive days. These wetlands experienced consistent diurnal environmental fluctuations between dawn and dusk, especially in terms of dissolved oxygen concentrations. Microbial community structure (16S rRNA gene sequencing) and active community structure (16S rRNA DGGE profiling) were analyzed with relation to diel fluxes (samples taken at dusk and dawn) within the water columns of wetlands.

## METHODS

### Sampling design

In August of 2015, four wetland sites were selected to explore the presence of diel fluxes of environmental conditions, as well as the microbial communities that may correspond to environmental fluctuations. Two of the wetlands, Main Marsh (MM) and North Marsh (NM) wetlands, were located on Beaver Island, MI, while the other two, Chipp-A-Waters Park oxbow lake (CW) and Pac Man Pond (PM) wetlands, were located in Mount Pleasant, MI. MM and NM were both part of the Miller’s Marsh system, however, the two branches of the marsh are separated by a Maple-Beech forest stand (Rowe, 2003). In the summer of 2015, samples were taken close to shore from two zones, nearshore (“A”; ~ 10 cm water depth), as well as out further into the wetland (“B”; ~ 30 cm water depth) within each site. Three replicate points were sampled in each zone. The two zones were established to examine whether community response to environmental diel fluxes was ubiquitous throughout the wetland or dependent upon water depth. Samples at each location were first taken at sunset, followed by dawn on the subsequent day. These times were chosen as O_2_ levels were likely to be higher at sunset (after photosynthesis occurred throughout the day) and respectively lower at dawn (where O_2_ could be depleted due to respiration and lack of photosynthesis at night). Collection was repeated in this fashion for two consecutive days at each location, totaling four sampling times at each wetland. Physical data were collected at each sampling location using a YSI (Yellow Springs, OH) hydrolab probe for measurement of O_2_ (DO), pH, conductivity (spCond), temperature (Temp), turbidity (Turbid), oxidation-reduction potential (ORP), and chlorophyll-a (Chl).

Water samples (both for DNA and chemical analysis) were collected by submerging one 500 mL bottle underwater to the depth just above the sediment/water interface, where the lid was opened and sealed once the bottle was filled. Of this water sample, 240 mL were immediately filtered through two consecutive sterile syringe push filters for DNA collection: a primary 2.7 μm filter, followed by a secondary 0.22 μm filter. The filters were immediately placed in sterile conical tubes, flash frozen in an ethanol and dry ice bath, and transported to the lab on dry ice. The first 30 mL of water to pass through the filters were collected into a glass vial, which contained HCl to reduce pH to 3 or lower for DOC measurements. Leftover unfiltered water for each sample was retained and transported to the lab on ice, where the water was pre-filtered through a 1.2 μm filter if needed, and then vacuum filtered through a 0.45 μm filter for nutrient analysis. DNA samples were stored at −80°C, DOC samples were stored at 4°C, and nutrient analysis samples were stored at −20°C.

In August of 2016, additional samples were collected from MM and NM wetlands for further analysis of RNA, as they experienced more pronounced and consistent diel O_2_ fluctuation regimes than CW and PM wetlands. A similar experimental design was implemented as the previous year, however, only zone “B” samples were taken (~ 30 cm water depth) as these locations experienced more dramatic diel fluxes than zone “A” the previous year. Samples were taken at each wetland first before dawn, and in the evening before sundown for one day in a similar way as described for DNA collection (as describe above) with the following modifications: Two 500 mL bottles were submerged underwater and opened above the sediment-water interface at each sampling point. For each water sample, 120 mL of water were filtered through each syringe filter system for a total of 4 filters, and a total of 460 mL of water were filtered for each sample unless filters became clogged before 120 mL of filtrate were able to be collected. One control sample was filtered in the field using sterile nanopure water at each wetland location. Filters were flash frozen in an ethanol and dry ice bath in the same manner as DNA preservation methods described above, as previous studies indicate that the use of RNA preservatives can result in a biased interpretation of bacterial community composition from freshwater samples (McCarthy et al., 2015). Filters were transported on dry ice and stored at - 80°C.

### Chemical analysis

DOC analysis was accomplished for the 2015 water samples using a Shimadzu (Kyoto, Japan) TOC-VCPH Total Organic Carbon Analyzer. For both 2015 and 2016 water samples, total N (TN), total P (TP), NH_3_, NO_3_^-^, and soluble reactive phosphorous (SRP) values for each sample were obtained through use of a Seal Analytical (Mequon, Wisconsin, USA) Quattro Bran+Luebbe Analyzer with a XY-2 sampler. Supplemental Table 1 contains raw chemical and physical data.

### Microbial community analysis

#### Microbial community rRNA gene sequencing

DNA was extracted from filters from the 2015 sampling effort using MoBio PowerSoil DNA extraction kits (Mo Bio, Carlsbad, CA). The quantity of the extracted environmental DNA for library preparation was assessed using a Qubit^®^ 2.0 fluorometer (Life Technologies, Carlsbad, CA). DNA samples were sent to Michigan State University for sequence library preparation and sequencing at the Research Technology Support Facility (Lansing, MI). The V4 region of the 16S rRNA gene was targeted in PCR with previously developed and commonly used primers 16Sf-V4 (515f) and 16Sr-V4 (806r) and protocol (Caporaso et al., 2012; Kozich et al., 2013). Generated amplicons were sequenced on a MiSeq high-throughput sequencer (Illumina, San Diego, CA) using paired-end 250 bp sequencing format.

Sequence data were quality filtered and analyzed using mothur v 1.35.1 (Schloss et al., 2009) following the MiSeq SOP (found at https://www.mothur.org) with minor modifications (full workflow for this project can be found at github.com/diel_wetland_comm_str). Paired-end sequences were joined into contigs. Sequences with homopolymers > 8 bases and sequences either less than 251 bp or greater than 254 bp were removed. Sequences were aligned against the Silva (v. 119) V4 rRNA gene reference database (Quast et al., 2012), and sequences which did not align within the V4 region were eliminated from further analysis. Using UCHIME (Edgar et al., 2011), chimeric DNA was searched for and removed from the dataset. Sequences were classified employing the Ribosomal Database Project (training set v. 9; Cole et al., 2013) using a confidence threshold of 80%. After taxonomic classification, if a sequence was identified as originating from chloroplast, eukaryotic, mitochondrial, or unknown sources, it was eliminated. Remaining sequences were clustered into Operational Taxonomic Units (OTUs) at the 0.03 sequence similarity level using the opticlust algorithm. Sequencing reads can be found in the Sequence Read Archive (SRA) under accession number SRP151564.

#### Microbial community rRNA DGGE analysis

The 2016 RNA samples were examined with PCR-DGGE (denaturing gradient gel electrophoresis) to characterize the active dominant microbial community composition. RNA was extracted using MoBio PowerWater RNA Isolation Kit according to manufacturer’s protocol. Extracted RNA was cleaned and concentrated using Zymo (Irvine, CA) Clean & Concentrator kit and quantified using a Qubit^®^ 2.0 fluorometer. RNA was converted to cDNA using Applied Biosystems (Foster City, CA) High Capacity cDNA Reverse Transcription kit. cDNA samples were amplified using bacterial primers 338f (ACT CCT ACG GGA GCG AGC AG) with a GC clamp attached to the 5’ end, and 519r (ATT ACC GCG GCT GCT GG) (Morgan et al., 2002). A standard PCR mix was used (Thermo Scientific, Waltham, MA) to which additional MgCl_2_ (Promega, Madison, WI) was added to bring the final concentration to 3.5 mM. In addition, Bovine Serum Albumin (Promega, Madison, WI) was added to a final concentration of 0.5 μg/μL. After an initial denaturation at 94°C for 5 min, 28 cycles of (1) denaturation at 92°C for 30 s, (2) annealing at 57°C for 20 s, and (3) extension at 72°C for 30 s were performed and followed by a final extension at 72°C for 7 min.

After PCR, DGGE was performed using 8% (w/v) polyacrylamide gels (37.5:1 acrylamide/bisacrylamide) with denaturing gradients that ranged from 30% to 52.5% (100% denaturant contains 40% [vol/vol] formamide and 7 M urea). Aliquots of 20 μL from the PCR reaction were subjected to DGGE. Each gel was run at 60 °C and 200 V for 330 min. After electrophoresis, gels were stained with GelGreen Nucleic Acid Stain (Biotium) and imaged using a ChemiDoc Touch Imaging System (Bio-Rad Laboratories, Hercules, CA). Bands on DGGE gels are assumed to represent members of the microbial community that make up > 1% of the population in a sample (Muyzer et al. 1993).

### Statistical analyses

The R statistical environment was used for statistical analyses (R Core Team, 2015). Principal Component Analysis (PCA) was used to visualize physical and chemical differences among samples. To address collinearity, highly correlated environmental variables (r > 0.7; p < 0.001) were removed from analysis save for one of the correlated variables in order to avoid exaggeration of PCA structure (DOC correlated with OrP, NH_3_^−^ correlated with TN, SRP correlated with TP). Permutational Multivariate Analysis of Variance (perMANOVA) (Anderson, 2001) was used to determine significant differences in physicochemical profiles among wetlands, sampling zones, and time of day.

One sample (out of three) of PM zone “B” at the first dusk sampling timepoint was removed from analysis as it was likely an artefactual result according to microbial community structure (95% composed of *Firmicutes* and *Actinobacteria*), which was distinct from replicates and all other samples (Supplemental Fig. 1). If a sample was represented by less than 1,000 sequences and Good’s coverage was < 90% (prior to removal of singletons and doubletons), the sample was also removed from further analyses. Good’s coverage estimates ranged from 94.2 - 99.6% among samples save for one sample which was estimated at 49.0% coverage due to low sequencing depth (n = 288 sequences), and was therefore, removed from further analyses. Variability in microbial community beta diversity among wetlands was visualized through NMDS based on Bray-Curtis dissimilarity among samples. The *envfit* function and a ranked Mantel test (where Euclidean distance was used to generate a distance matrix based on geochemical profiles) from the Vegan package (Oksanen et al., 2007) were used to determine relationships between microbial community structure and environmental variables among wetlands. perMANOVA was used to explore whether wetlands were distinct in microbial community structure, and whether interactions between wetland, sampling zone, and time of day were also significant in explaining differences in microbial community profiles.

To further assess the degree of variability among microbial communities between diel sampling points, each wetland was analyzed individually to control for differences in community structure among wetlands. NMDS was performed for each wetland, as well as for each zone of sampling within each wetland. To test for significant differences between time and sampling points, perMANOVA was performed for each wetland and zone of sampling within each wetland.

DGGE profiles of different samples within each wetland were compared by calculating Jaccard distance coefficients based on the presence/absence of bands. Significant differences between time points were analyzed by performing perMANOVA to test for significant differences in active communities between dawn and dusk time points.

To explore whether relationships between microbial subnetworks, taxa, and O_2_, pH, or temperature fluxes existed within each wetland, weighted correlation network analysis (WGCNA) was implemented using the *WGCNA* package (Langfelder & Horvath, 2008; Langfelder & Horvath, 2012) as previously described (Guidi et al., 2016; Henson et al., 2016; Horton et al., *in review*) with minor modifications. In summary, WGCNA analysis was applied to each sampling zone (A versus B) within each wetland separately, totaling 8 separate analyses. An OTU was removed from further analysis if it did not appear at least twice within at least 25% of the samples explored to control for erroneous correlations (similar to as applied by Henson et al., 2018). OTU abundances were normalized using variance stabilizing transformation (VST). Constructed dissimilarity matrices were then raised to a soft threshold power to ensure scale-free topology. This power was chosen on an individual basis for each dissimilarity matrix based on which soft threshold power met the assumption of scale-free topology, while allowing for the greatest connectivity among OTUs within the network. Topological overlap matrices (TOMs) were created, and subnetworks were generated using TOMs and hierarchical clustering. Pearson correlations were calculated between the assigned eigenvalue for each subnetwork (defined as the first principal component representing a given subnetwork) and dissolved oxygen, pH, and temperature levels to explore relationships between subnetworks of microbial taxa and diel fluxes. The subnetworks with the strongest correlations (if r > 0.7, p < 0.01) to environmental variables were selected for further exploration. Partial least squares regression (PLS) models were employed to explore the ability of subnetworks to predict levels of O_2_, where leave-one-out cross-validation (LOOCV) predicted values were used to test the ability of the PLS model to predict measured values. Variable importance in projection (VIP) (Chong & Jun, 2005) defined the contribution of each OTU in predicting values of environmental variables during PLS. For visualization purposes, the minimum correlation (r) between two OTUs to constitute a relationship was delineated between 0.1 and 0.15 dependent upon the strength of OTU - OTU correlations within each subnetwork. The sum of relationships an OTU possesses with other OTUs within a network was defined as “node centrality”.

## RESULTS AND DISCUSSION

### Each wetland experienced distinct diel fluxes in physical and chemical conditions

The four tested wetlands were distinct in physical and chemical conditions according to perMANOVA (r^2^ = 0.435, p < 0.001) and pairwise comparisons (Supplemental Table 2). Specifically, Miller’s Marsh wetland sites (MM and NM) exhibited higher OrP, temperature, average O_2_ levels, and pH than wetlands sampled in Mount Pleasant, MI (CW and PM) (Fig. 1; Supplemental Table 1). CW and PM wetlands were relatively higher in specific conductivity, turbidity, and chlorophyll-a concentrations. A beta-dispersion test (Anderson, 2006; Anderson et al., 2006) found that physicochemical variance among samples within each wetland was not significantly different among wetlands. Further, perMANOVA found significant interaction effects among wetlands, sampling zones, and time of day (Supplemental Table 3), suggesting that diel physicochemical fluxes were variable among wetlands and sampling zones. Wetlands located on Beaver Island, MI (MM and NM) experienced relatively steeper and more consistent diel fluctuations in dissolved oxygen than those located within Mount Pleasant, MI (CW and PM), particularly within sampling zone “B” (Fig. 2). The degree to which temperature and pH varied between dawn and dusk time points were relatively consistent among all wetlands, save for relatively larger variability in pH within site MM.

**Figure 1.**
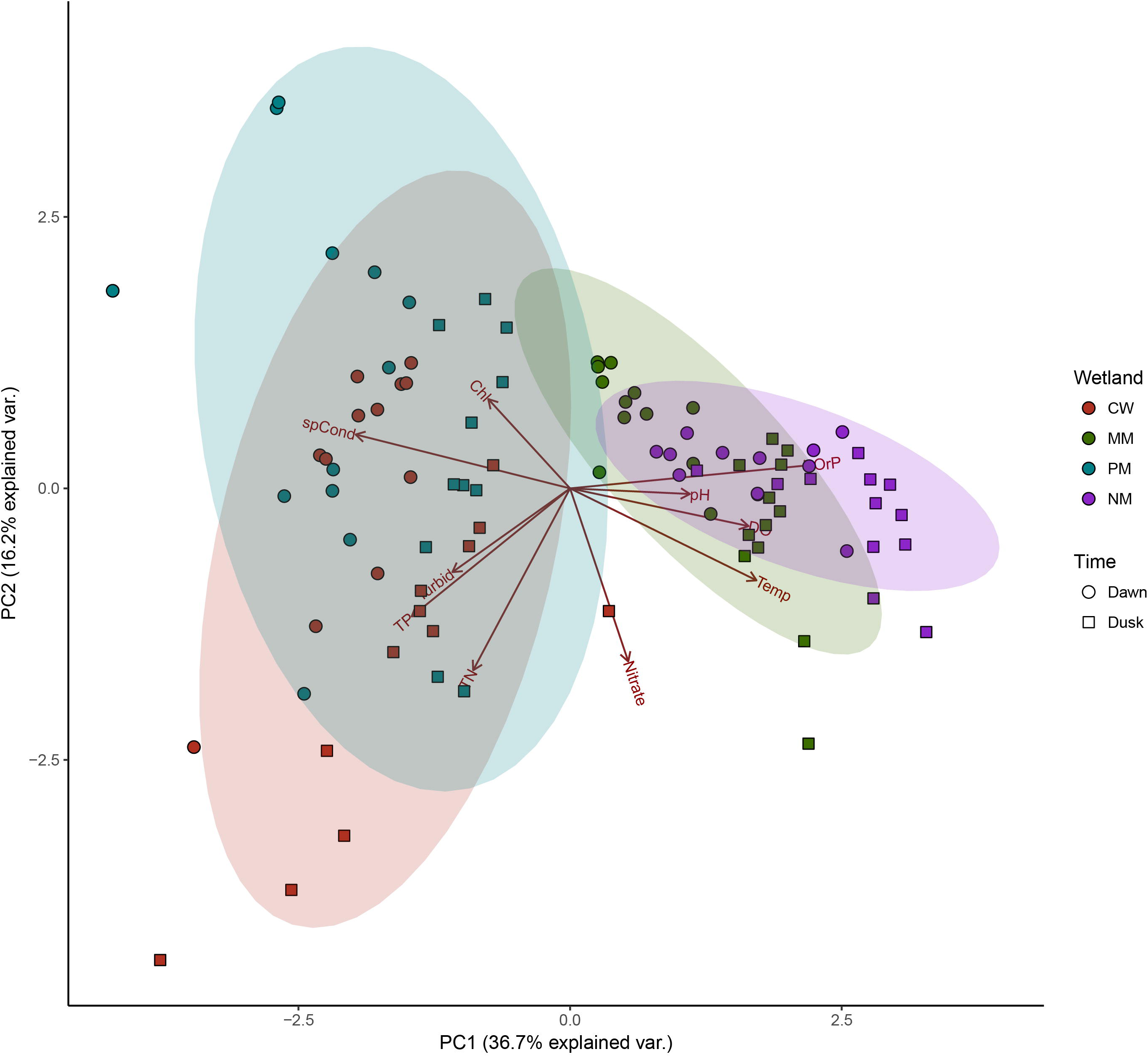
Principal component analysis of samples characterized by geochemical signatures among wetlands (colors) and time points (shapes). Principal component 1 (36.7% explanatory) and principal component 2 (16.2% explanatory) are displayed. Vectors represent individual geochemical variables separating samples in two-dimensional space. Ellipses represent 95% confidence intervals.

**Figure 2.**
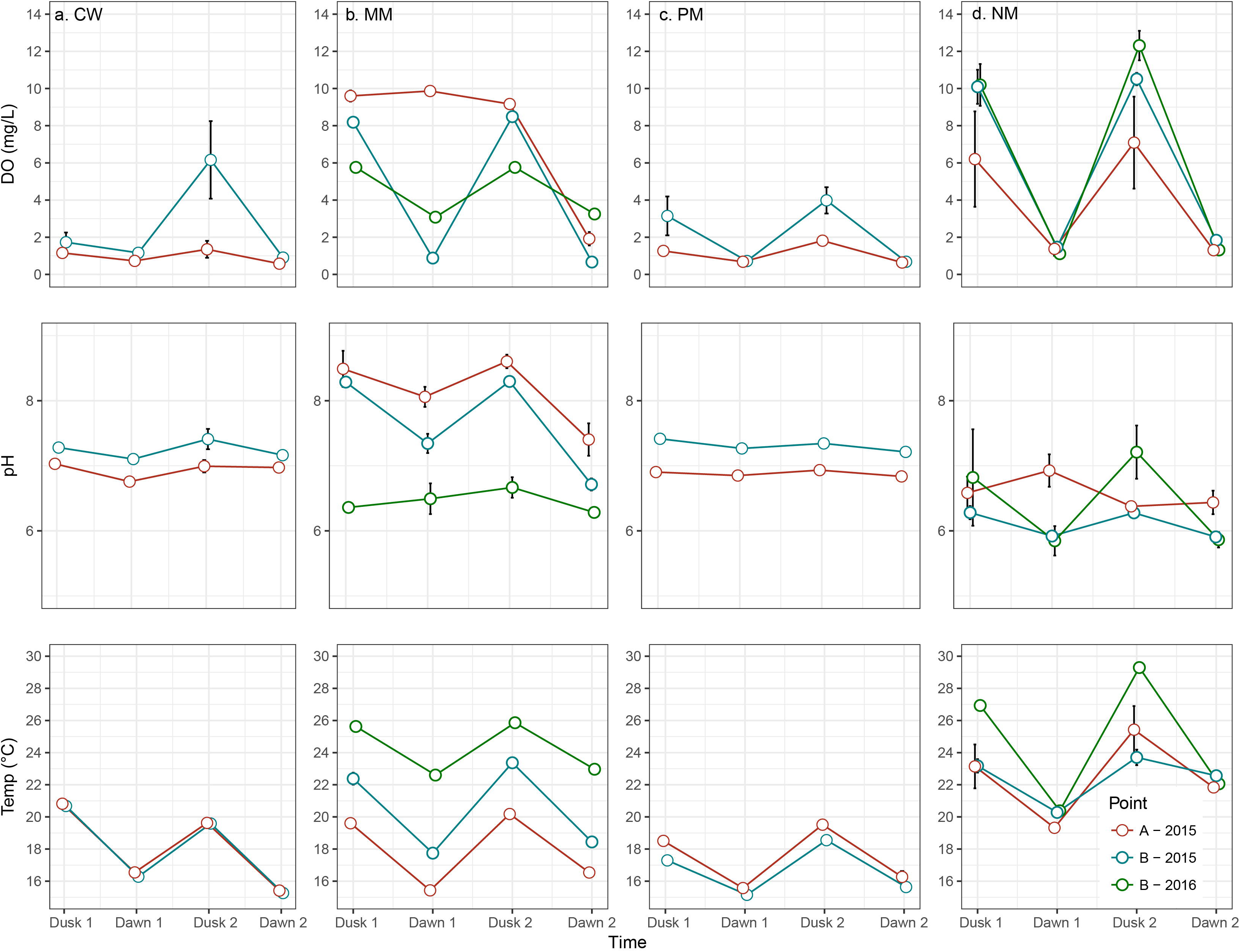
Diel fluctuation of dissolved oxygen (DO), pH, and temperature (Temp) for wetlands a.) CW, b.) MM, c.) PM, and d.) NM during 2015 and 2016. Line color represents sampling zone and year sampled consistent with the legend. Error bars represent +/- standard error.

### Total microbial community structure was primarily stable throughout diel fluxes

After quality filtering of sequence data, a total of 11,655,362 sequences remained. Both perMANOVA (r^2^ = 0.516; p < 0.001) and pairwise perMANOVAs demonstrated that microbial communities were unique to each wetland (Fig. 3; Supplemental Table 2). The differences among microbial communities were likely driven by distinct environmental conditions among each wetland, as wetland physicochemistry and community structure were related according to a ranked Mantel test (r = 0.577, p ≤ 0.001). Corroborating this, the NMDS ordination was correlated to differences in multiple environmental variables (Fig. 3). Wetlands with distinct microbial community structure are frequently distinct in geochemical properties (Peralta et al., 2013; Ligi et al., 2014), which is consistent with microbial community patterns in the wetlands studied here. As microbial communities were distinct among all wetlands, individual wetlands were independently examined for relationships between microbial community structure and environmental fluxes. Further, as interaction effects were significant between wetland and sampling zone (perMANOVA, r^2^ = 0.06, p ≤ 0.001), each wetland was also explored to examine whether microbial communities were spatially distinct.

**Figure 3.**
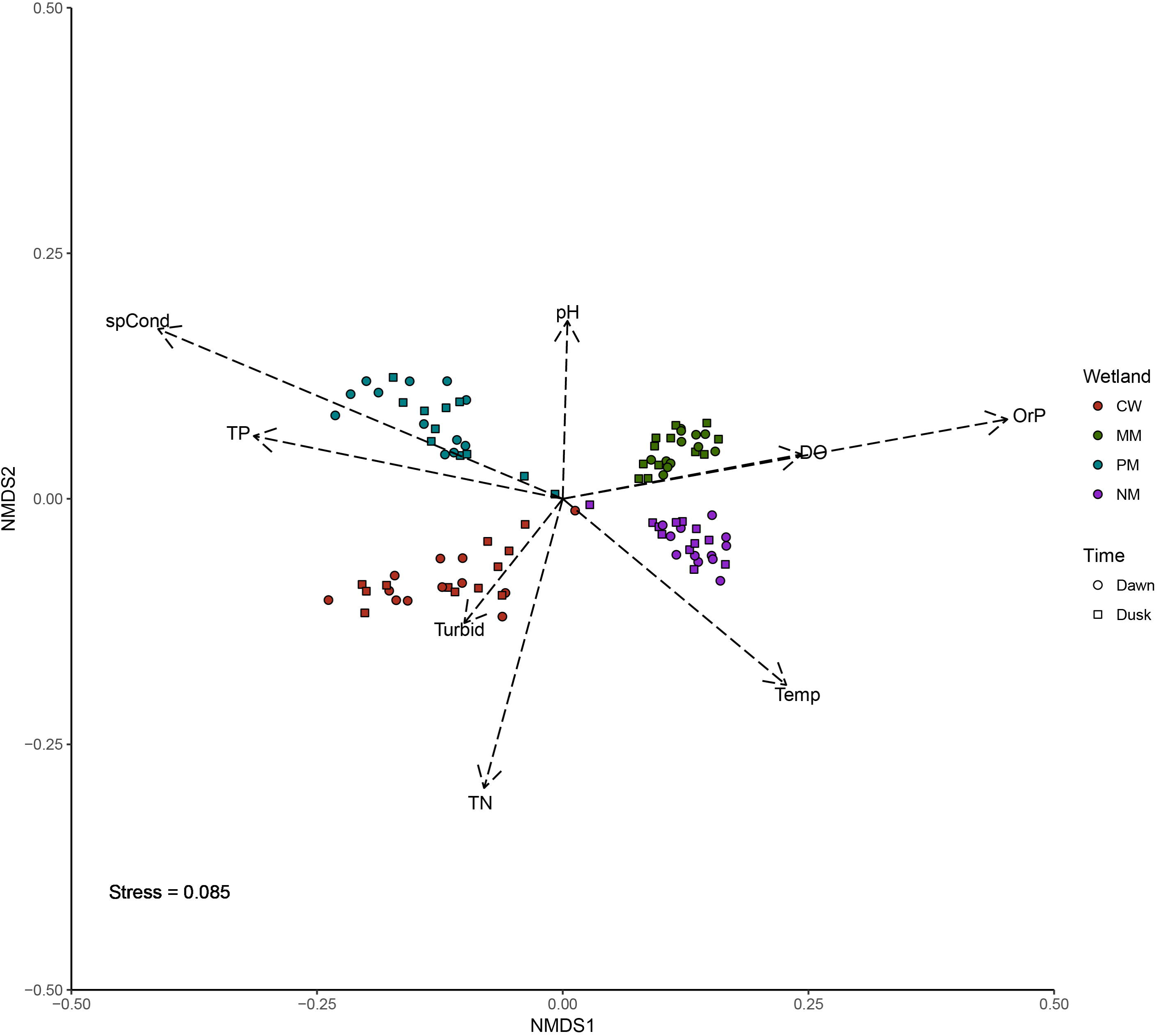
Nonmetric multidimensional scaling separating samples based upon microbial community structure. Colors represent wetland, while shapes represent time point. Vectors represent significant correlations (p < 0.01) of environmental variables to NMDS structure, and vector length represents strength of correlation.

Microbial community structure consistently varied between sampling zones according to perMANOVA and NMDS within each wetland (Fig. 4; Supplemental Table 4). These results show that spatial differences exist within the tested 3-meter scale and underscore the importance of analyzing spatially distinct zones within wetland systems, as microbial communities can be distinct in community structure throughout a wetland, and therefore, may respond to diel fluxes differentially. Similar types of spatial variability in bacterial communities has also been found in other studies which explored wetland microbial communities (Song et al., 2012; Narrowe et al., 2017). While microbial community structure can also be impacted by dominant vegetative type (Tang et al., 2011), the samples collected in this study were taken within the same vegetation type (defined as at least 75% of one morphotype). Thus, it is reasonable to suggest that point of sampling has a profound effect on microbial community structure even at finer spatial scales than “vegetation zone” within wetlands, possibly related to differences in water depth.

**Figure 4.**
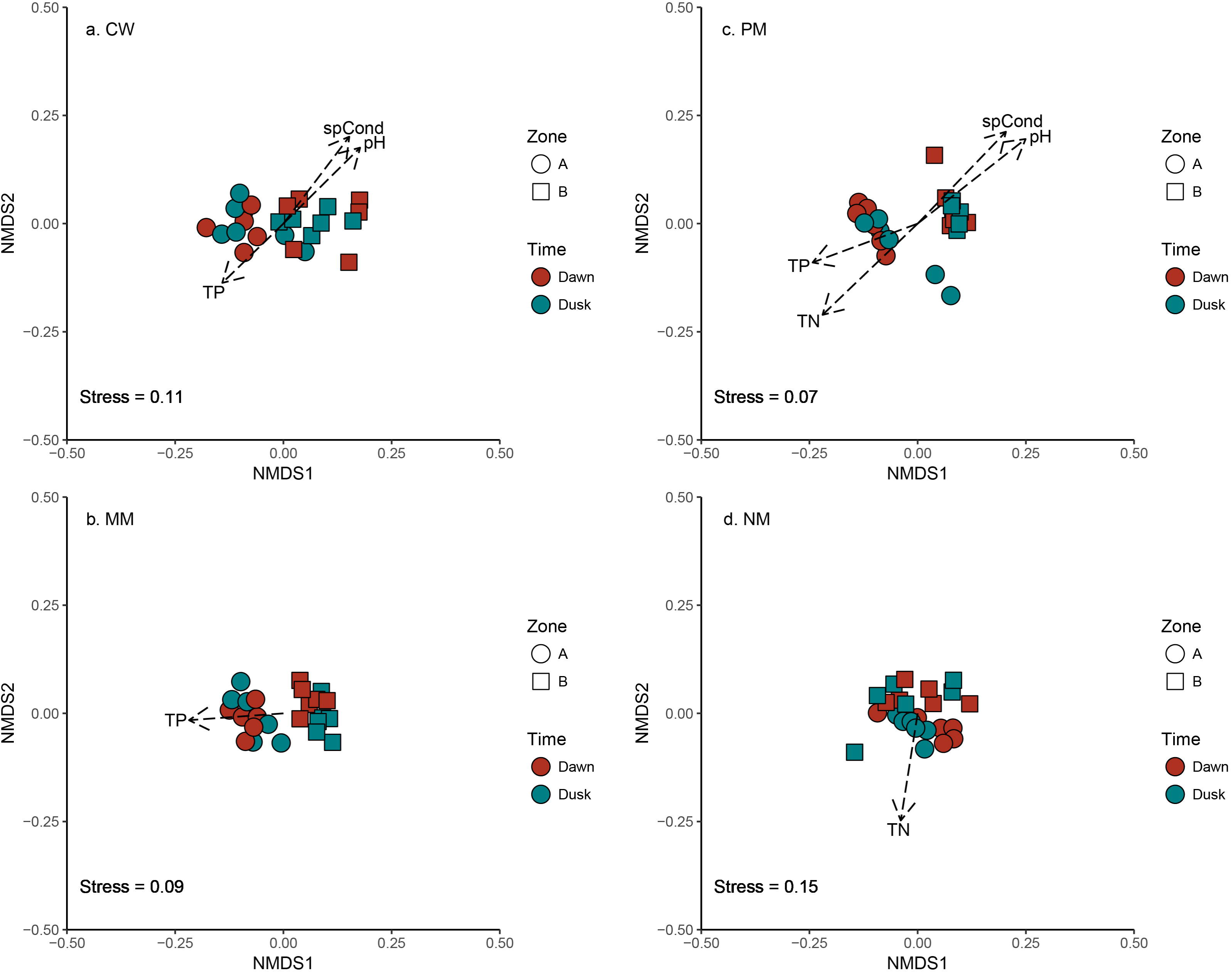
NMDS examining microbial community structural relationships among samples within individual wetlands a.) CW, b.) MM, c.) PM, and d.) NM. Colors represent sampling time point while shapes represent zone of sampling. Vectors represent significant correlations (p < 0.01) of environmental variables to NMDS structure, and vector length represents strength of correlation.

Changes in microbial community beta diversity between dawn and dusk time points were only significant in zone “B” of MM according to perMANOVA (r^2^ = 0.140, p = 0.007). Interestingly, total community structure was variable in zone “B” of MM (based on sequencing
data from DNA), but DGGE analysis of active microbial community structure (via RNA extractions) did not oscillate between night and day time points within either MM or NM (Fig. 5). These data suggested that the dominantly active microbial community may not be driving daily shifts in microbial community structure found in MM, and that a subset of less abundant microbial taxa could be driving community-level changes in MM. Overall, these results contrast with those found by Kearns et al. (2017), where it was established that the active microbial community shifted over a daily cycle within salt marsh water. However, salt marshes are unique in that they are transient systems which alternate between influxes of saltwater and periods of stagnation, whereas the freshwater wetlands studied here were closed, stable systems.

**Figure 5.**
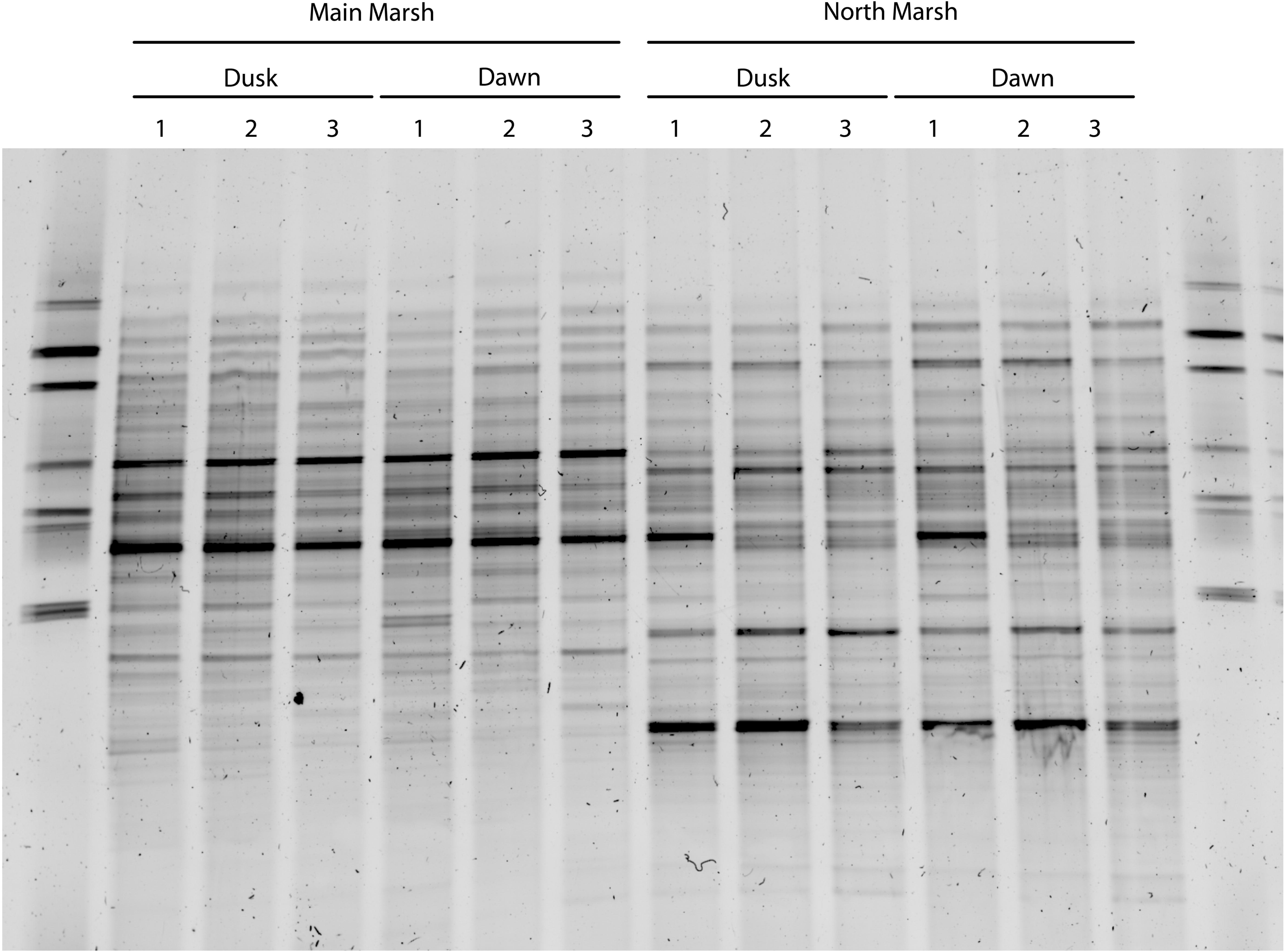
DGGE profiles of RNA samples taken in 2016 and MM and NM wetlands during dawn and dusk time periods. Numbers represent replicate samples. The two unlabeled lanes were loaded with a marker produced in our laboratory from the 16SrDNA of several environmental isolates.

Dominant microbial community members within closed freshwater wetland water columns may be adapted to diurnal fluctuations of environmental variables (such as O_2_, temperature, and pH). It has been shown that some microbial taxa may retain the ability of functional plasticity in the face of disturbance (Shade et al., 2012), and dominant microbial community members may exhibit such plasticity through adaptation to daily fluctuation regimes. Microbial dormancy may also be contributing to a lack of community structure differences between dawn and dusk within wetland water. Inactive community members have been found to constitute close to 30% of communities within freshwater systems (Lennon & Jones, 2011) and estimated at up to 62% of community membership in a saltwater marsh which experienced diel fluxes (Kearns et al., 2017). Further, DNA within freshwater environments can remain detectable for several days after removal of the DNA source (Dejean et al., 2011), thus persistence of microbial community DNA from dead cells may contribute to masking fine-scale community composition shifts.

### Microbial subcommunities respond to diel O_2_ fluxes in freshwater wetlands

Diel fluxes showed little to no influence on beta diversity, however, these communities were further explored at a finer resolution to determine if potential subnetworks of taxa may have been impacted by fluctuating environmental conditions. Network analysis, via WGCNA, showed that each wetland harbored unique subnetworks of taxa that correlated with fluctuating environmental factors. Specifically, two subnetworks correlated with fluctuating dissolved oxygen levels within wetlands MM and PM.

Within MM sampling zone “B” (which experienced steep oxygen fluctuation regimes), the subnetwork most related to dissolved oxygen concentrations was 34.6% predictive of dissolved oxygen levels according to PLS modeling. Additionally, the same subnetwork that correlated to DO (r = 0.7, p = 0.01) also significantly correlated with shifts in temperature (r = 0.7, p = 0.01) (Fig. 6). This subnetwork was composed of taxa spanning several phyla and individual OTUs whose relative abundances were (either positively or negatively) correlated with dissolved oxygen levels. OTUs related to *Alphaproteobacteria* possessed some of the highest VIP scores and were positively related to DO concentrations, along with other OTUs associated with *Mycobacterium* (*Actinobacteria*), *Sphingobacteriales* (*Bacteroidetes*), and *Armatimonadetes* Gp1. Interestingly, an aerobic representative within Gp1 of *Armatimonadetes* (*Armatimonas rosa*) was isolated from the rhizoplane of a common wetland grass *Phragmites australis* and has also been shown to be incapable of nitrate respiration or fermentation (Tamaki et al., 2011). Therefore, this isolate may be sensitive to diel O_2_ fluxes, which have been shown to occur within wetland plant rhizospheres (Nikolausz et al., 2008). OTUs which possessed high VIP scores and were negatively correlated to dissolved oxygen included those related to *Methylococcales* and other *Proteobacteria*, as well as *Verrucomicrobia* Subdivision 3. *Methylococcales*, an order of bacteria representing known methanotrophs, has been shown to decrease in abundance with increasing O_2_ levels within the oxygen minimum zone (OMZ) of Golfo Dulce, Costa Rica (Padilla et al., 2017). Further, *Methylococcales* have been found to pair oxidation of methane with partial denitrification (Padilla et al., 2017). Thus, the relative abundances of *Methylococcales* OTUs documented in this study may be responding to fluctuations in diel chemical and physical conditions.

**Figure 6.**
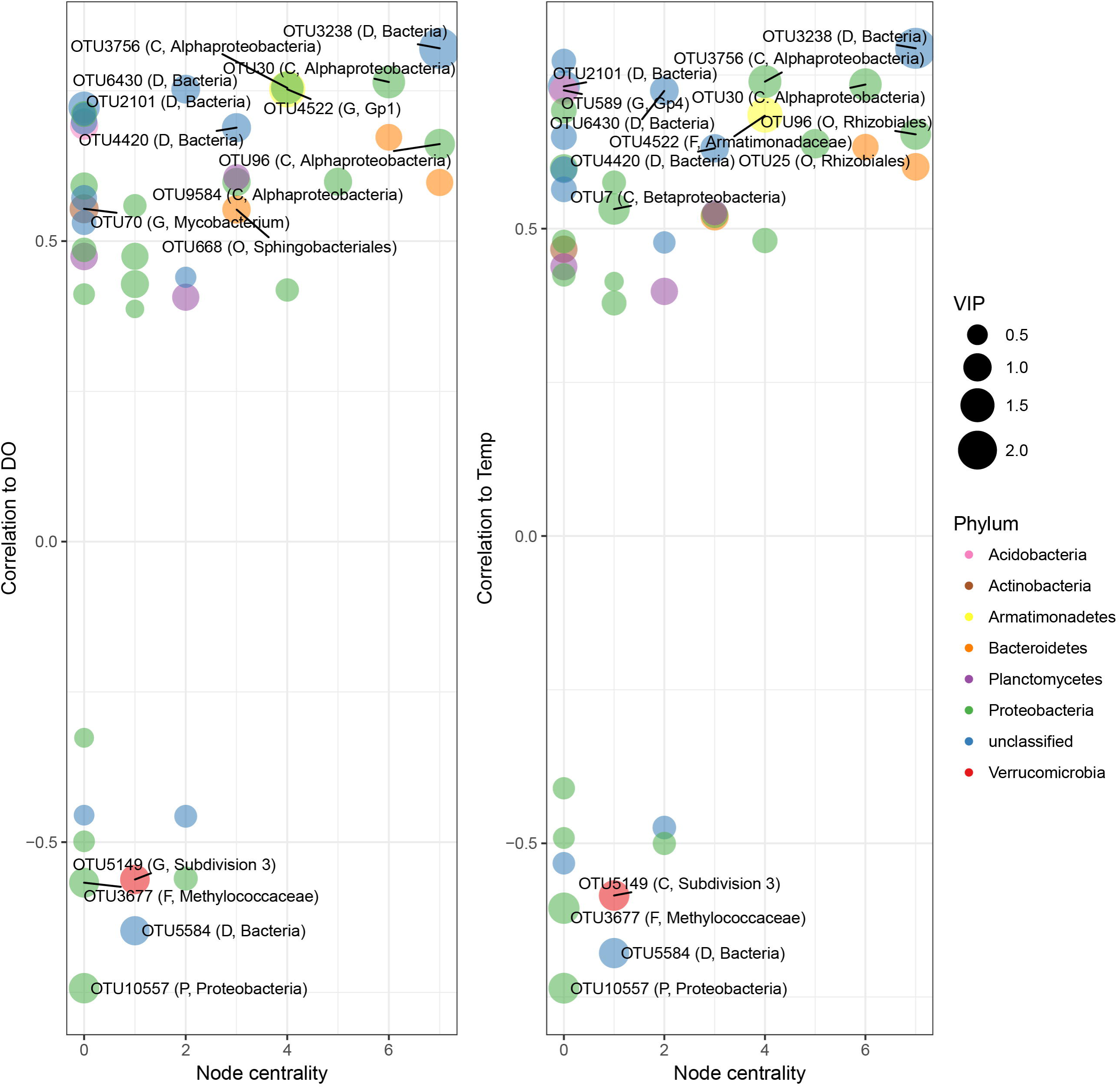
Subnetworks with significant relationships to a.) DO concentrations and b.) temperature from zone “B” of MM. Points correspond to OTUs with network membership. Color of node corresponds to Phylum, and shape of the node represents relative VIP score. Correlation to environmental variables for individual OTUs rests on the y-axis, and “node centrality” rests on the x-axis. OTUs with the top 15 VIP scores are labeled, with corresponding lowest taxonomic resolution at which they are identified followed by taxonomic identification (D: Domain; P: Phylum; C: Class; O: Order; F: Family; G: Genus).

While there were no significant differences in total microbial community structure between dawn and dusk within PM, WGCNA found a subcommunity that was significantly related to the narrow range of dissolved oxygen fluctuations within this wetland. This subcommunity was 66.8% predictive of dissolved oxygen concentrations in zone “B” of PM according to PLS modeling. This subnetwork correlated with DO (r = 0.78, p = 0.004), and also correlated with pH (r = 0.83, p = 0.002) (Fig. 7). OTUs with high VIP values that positively correlated with dissolved oxygen included *Aquabacterium (Betaproteobacteria), Rheinheimera (Gammaproteobacteria), Bacteroidetes*, and *Verrucomicrobia* Subdivision 3. *Aquabacterium* and *Rheinheimera* have been isolated from freshwater sources and characterized as facultative anaerobes capable of nitrate reduction (Kalmbach et al., 1999; Merchant et al., 2007; Chen et al., 2010). Conversely, OTUs that negatively correlated with dissolved oxygen were *Acidobacteria* Gp 6, *Chloroflexi*, and candidate phylum OD1 (also known as *Parcubacteria*). Interestingly, *Parcubacteria* have been suggested as taxa with reduced metabolic capabilities, potentially due to obligate parasitic strategies according to genomic analyses (Nelson & Stegen, 2015), and as such, may be dependent on other organisms which are sensitive to diel fluxes. It is also possible that the two taxonomically distinct subnetworks found to be related to O_2_ within MM and PM contained some microbial taxa that maintain similar metabolic strategies which coincide with redox-controlled elemental transformations that commonly occur in wetlands (Jørgensen et al., 1979; Laursen & Seitzinger, 2004; Harrison et al., 2005). However, more research would be necessary to understand whether functional redundancy of microbial communities existed among these wetlands.

**Figure 7.**
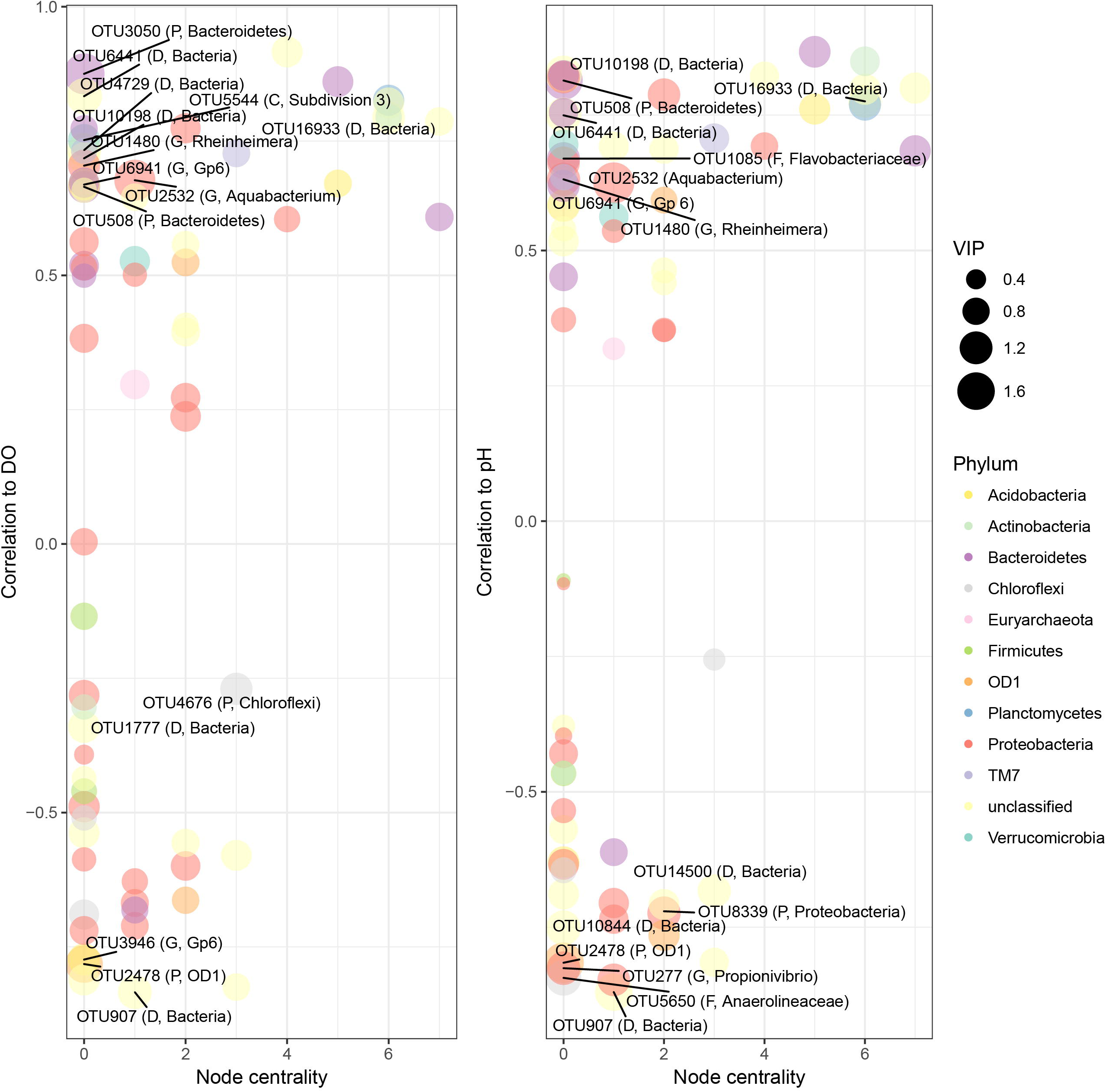
Subnetworks with significant relationships to a.) DO concentrations and b.) pH from zone “B” of PM. Points correspond to OTUs with network membership. Color of node corresponds to Phylum, and shape of the node represents relative VIP score. Correlation to environmental variables for individual OTUs rests on the y-axis, and “node centrality” rests on the x-axis. OTUs with the top 15 VIP scores are labeled, with corresponding lowest taxonomic resolution at which they are identified followed by taxonomic identification (D: Domain; P: Phylum; C: Class; O: Order; F: Family; G: Genus).

Interestingly, NM experienced the most dramatic daily O_2_ fluxes, and did not possess any unique subnetworks of microbial taxa related to oxygen fluxes as found in MM within the same wetland system. These data further allude to idiosyncrasy that may exist among the microbial community response to diel O_2_ fluxes within wetlands, possibly dependent upon microbial community taxonomic membership and physicochemical differences among wetlands.

## CONCLUSIONS

It is evident that the wetlands examined in this study were unique in both physicochemical and microbial fingerprints, and specific elements of these wetland ecosystems may have influenced the degree to which the microbial community structure responded to natural diel fluxes. Broad community beta diversity patterns were found to significantly differ between dawn and dusk time periods in only one out of eight wetland zones. Rather, small subnetworks of taxa were more often found to shift with oxygen levels within each wetland. Therefore, it is likely that dominant microbial taxa within these freshwater wetlands (and wetland zones) remain structurally stable throughout the day, while smaller subsets of community members are more sensitive to daily environmental fluxes. Further research is necessary to fully understand how diel fluxes impact the function of microbial communities. Nevertheless, this research highlights the importance of exploring microbial communities at finer resolutions (subcommunities) which may be masked by examining patterns across the entire microbial community.

## ACKNOWLEDGEMENTS

We would like to thank Miranda Hengy, Adam Byrne, and Shanker Tamang for their assistance in the sampling of these wetlands and John Gordon and the CMU Biological Station staff for all their logistical assistance. Funding for this project was provided by the CMU Institute for Great Lakes Research. This is contribution number XXX of the Institute for Great Lakes Research.

## SUPPLEMENTAL INFORMATION

Supplemental information, figures, and tables can be found online at the location https://github.com/horto2dj/diel_wetland_comm_str.

